# MMP2 As An Independent Prognostic Stratifier In Oral Cavity Cancers

**DOI:** 10.1101/723650

**Authors:** Caroline Hoffmann, Sophie Vacher, Philémon Sirven, Charlotte Lecerf, Lucile Massenet, Aurélie Moreira, Aurore Surun, Anne Schnitzler, Jerzy Klijanienko, Odette Mariani, Emmanuelle Jeannot, Nathalie Badois, Maria Lesnik, Olivier Choussy, Christophe Le Tourneau, Maude Guillot-Delost, Maud Kamal, Ivan Bieche, Vassili Soumelis

**Affiliations:** Paris-Saclay University, Paris, France; INSERM U932 research unit, Immunity and Cancer, Paris, France; Department of Surgical Oncology, Institut Curie, Paris & Saint-Cloud, France; Department of Genetics, Institut Curie, Paris, France; Department of Drug Development and Innovation (D3i), Institut Curie, Paris & Saint-Cloud, France; SIREDO Cancer Center (Care, innovation and research in pediatric, adolescents and young adults oncology), Institut Curie, Paris, France; Paris Descartes University, Sorbonne Paris Cité, Paris, France; Department of pathology, Institut Curie, Paris, France; Biological Resources Center, Institut Curie, Paris, France; INSERM U900 research unit, Saint-Cloud, France; Center of Clinical Investigation, CIC IGR-Curie 1428, Paris, France; INSERM U1016 research unit, Paris Descartes University, Faculty of Pharmaceutical and Biological Sciences, Paris, France; Clinical immunology department, Institut Curie, Paris, France. Current address of the author is Hospital St Louis, Immunology & Histocompatibility Laboratory, Paris, France

**Keywords:** Biomarker, Metalloproteinase, Prognosis, Secretome, Squamous cell carcinoma, Head and Neck, Oral cavity

## Abstract

**Background:** Around 25% of oral cavity squamous cell carcinoma (OCSCC) are not controlled by standard of care. Identifying those patients could offer them possibilities for intensified and personalized regimen. However, there is currently no validated biomarker for OCSCC patient selection in a pre-treatment setting.

**Patients and methods:** Our objectives were to determine a robust and independent predictive biomarker for disease related death in OCSCC treated with standard of care. Tumor and juxtatumor secretome were analyzed in a prospective discovery cohort of 37 OCSCC treated by primary surgery. Independent biomarker validation was performed by RTqPCR in a retrospective cohort of 145 patients with similar clinical features. An 18-gene signature (18G) predictive of the response to PD-1 blockade was evaluated in the same cohort..

**Results:** Among 29 deregulated molecules in a secretome analysis, we identified soluble MMP2 as a prognostic biomarker. In our validation cohort (n=145), high levels of *MMP2* and *CD276*, and low levels of *CXCL10* and *STAT1* mRNA were associated with poor prognosis in univariate analysis (Kaplan-Meier). *MMP2* (p = 0.001) and extra-nodal extension (ENE) (p = 0.006) were independent biomarkers of disease-specific survival (DSS) in multivariate analysis, and defined prognostic groups with 5-year DSS ranging from 36% (*MMP2*highENE+) to 88% (*MMP2*lowENE-). The expression of 18G was similar in the different prognostic groups, suggesting comparable responsiveness to anti-PD-1.

**Conclusion:** High levels of MMP2 was an independent and validated prognostic biomarker, which may be used to select poor prognosis patients for intensified neoadjuvant or adjuvant regimens.

## INTRODUCTION

Oral cavity squamous cell carcinoma (OCSSC) patients treated by primary surgery undergo post-operative surveillance, adjuvant radiotherapy, or chemo-radiotherapy, according to clinical and histopathological parameters that include disease stage, nodal involvement, extranodal extension (ENE), perineural invasion (PNI), vascular embols (VE) and resection margin status (1). Despite those numerous clinical decision parameters, around 25% of OCSCC will present an unpredictable early and/or severe recurrence (2), (3), (4). Even the local failures that are eligible to the best treatment option, that is salvage surgery (5), (6), (7), have a poor prognosis with a median overall survival ranging from 20 to 30 months (4), (8). Accurately identifying those high-risk patients would allow proposing them an intensified and risk-adjusted therapy, such as neoadjuvant chemotherapy or immunotherapy.

Neoadjuvant chemotherapy has failed to show benefit in head and neck squamous cell carcinoma (HNSCC), possibly because trials were made in unselected Stage III/IV HNSCC population (9), (10). Immunotherapy is a new treatment modality, and its interest as neoadjuvant treatment is currently being evaluated (11), (12), (13). Numerous prognostic markers have been proposed for OCSCC, but none of them has shown independent validation, and translation to clinical practice (14). In this study, we used a biology-driven exploratory strategy, in order to identify a robust predictive biomarker for early severe recurrence and disease related death in primary OCSCC after treatment by standard of care. We found MMP2 as fulfilling those criteria, and when combined to nodal involvement, providing a simple and efficient patient stratification scheme.

## RESULTS

### Human primary tumor secretome analysis identified 29 deregulated molecules

To identify candidate biomarkers, we chose an unbiased approach applied to human primary tumors, in order to ensure physiopathological relevance. We used a tumor explant-culture system to analyze the soluble microenvironment in a prospective discovery cohort of 37 OCSCC patients treated by primary surgery (Table S1). Fresh standardized tumor and juxtatumor (non-involved) specimens were cultured for 24h at 37°C, and we measured a panel of 49 soluble molecules relevant to multiple cancer pathways, such as immunity, chemotaxis, tumor growth, angiogenesis, and tissue remodeling. We identified 25 molecules increased, and 4 decreased, in the tumor tissue (Fig 1, Table S2). CXCL9, the metalloproteinases (MMP) MMP1, MMP2 and MMP9, plasminogen activator inhibitor (PAI-1) and resistin were among the molecules most increased in tumors, and MCP-1 (CCL2) in juxtatumors. SCF, multiple cytokines (IL-1b, TNFa, IL-15), growth factors (GM-CSF, VEGF) and chemokines (MDC, TARC) were also increased in the tumor, as compared to juxta-tumor samples (Fig 1). The cytokines IL-9, TNFb, TSLP, IL-21 were never detected (Fig 1). This provided a global, unbiased protein level profiling of the OCSCC tumor secretome.

**Fig 1.**
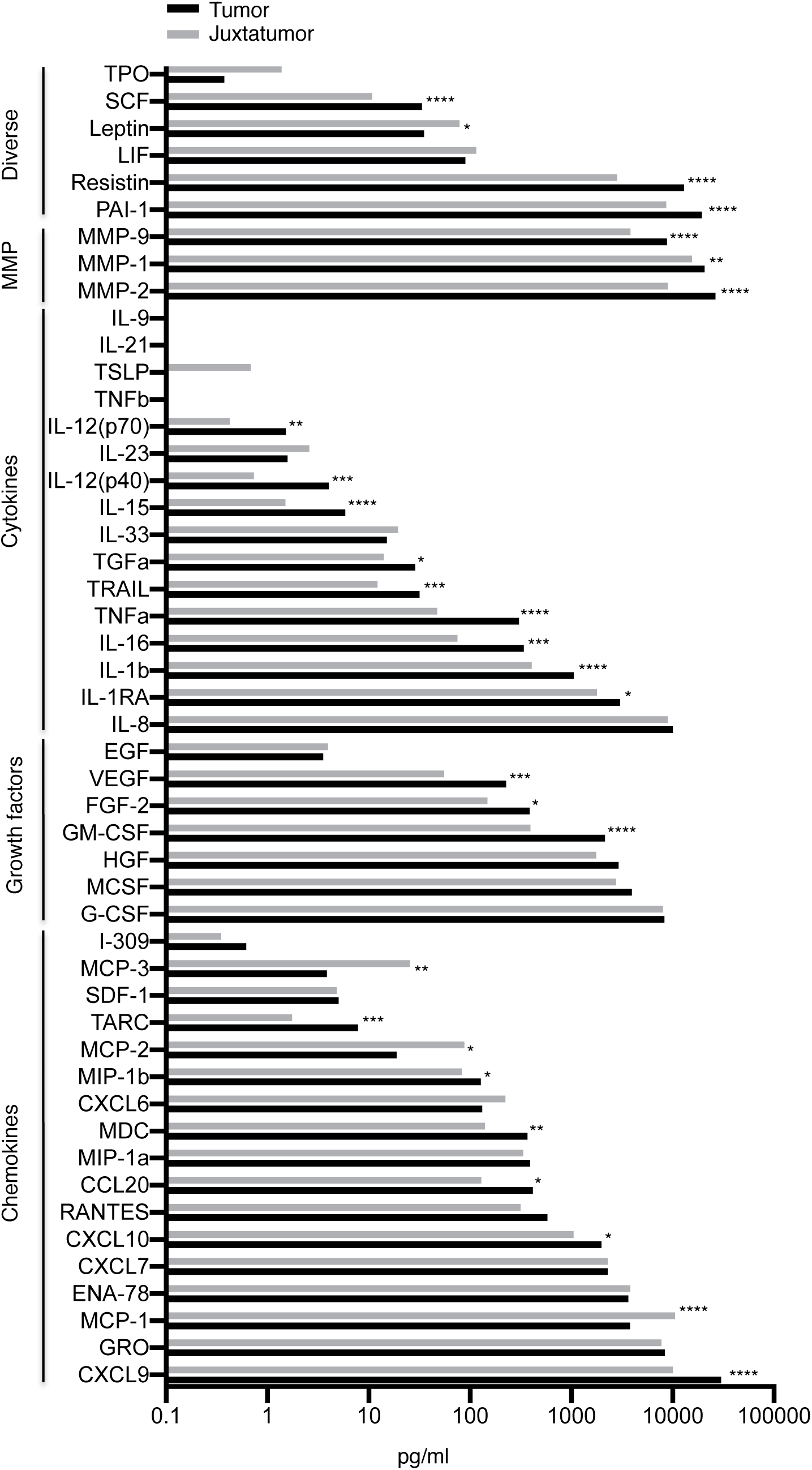
Tumor secretome analysis identified 29 deregulated molecules. Quantification of 49 molecules from the soluble microenvironment of 37 OCSCC and paired juxtatumor tissue. P-values obtained by Wilcoxon tests are represented by range: * < 0.05, ** < 0.01, *** < 0.001, **** < 0.0001.

### High levels of soluble MMP2 were associated with poor prognosis

Patients were classified as severe if they had a disease-specific survival (DSS) of less than 36 months and /or a disease-free survival (DFS) of less than 12 months, and could not achieve a second remission (unsuccessful salvage procedures and/or permanent palliative treatment). Among the 29 deregulated secretome molecules, analyzed as candidate biomarkers, MMP2 was the only molecule expressed at significant higher levels among severe patients as compared to non-severe (p = 0.007) (Table S3). ROC curve defined 29.3 ng/ml as the optimal cut-off for soluble MMP2, with a sensitivity of 100% and a specificity of 71.4 % to identify severe cases (Fig2A). MMP2high tumors were associated with reduced DSS (p = 0.001), overall survival (OS) (p = 0.012) and DFS (p = 0.003) (Fig 2B).

**Fig 2.**
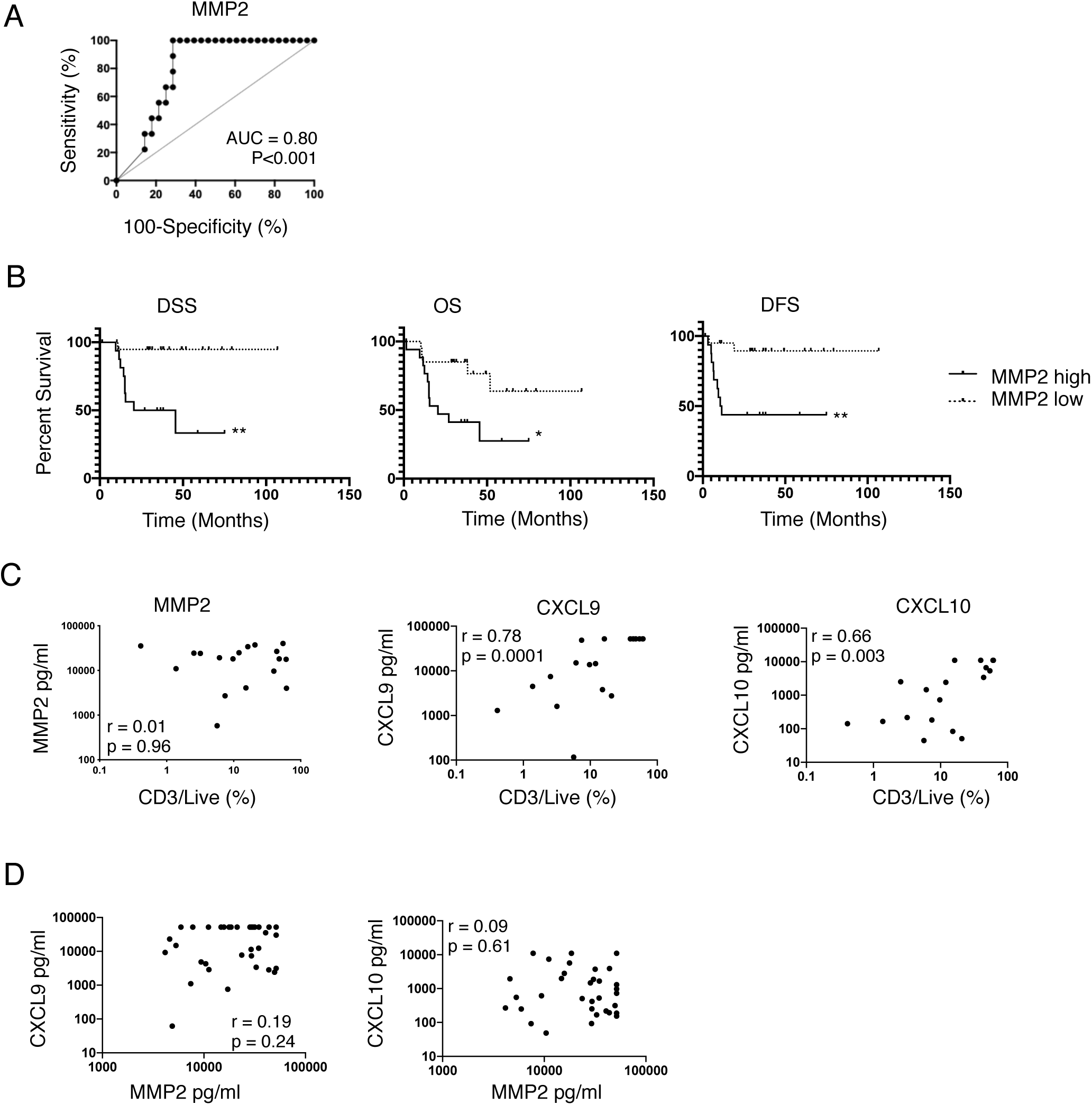
Soluble MMP2 is a prognostic biomarker of OCSCC, independent of T cell infiltration. A. ROC curve of soluble MMP2 for severity criteria (DSS < 36 months and /or a DFS < 12 months followed by permanent palliative treatment). The optimal threshold was 29.3 ng/ml. B. DDS, DFS and OS survival curves according to soluble MMP2 level, define as high or low relatively to the threshold defined in “B”. C. Correlation between CD3 in live cells and soluble MMP2 (left), CXCL9 (center) and CXCL10 (right), in tumors of 18 HNSCC patients. r values are Spearman correlation coefficients. D. Correlation between soluble MMP2 and CXCL9 (left) and CXCL10 (right), in 37 OCSCC samples. r values are Spearman correlation coefficients. Abbreviations. OCSCC: oral cavity squamous cell carcinoma, ROC: receiver operating characteristic, DSS: disease specific survival, DFS: disease free survival, OS: overall survival, HNSCC: head and neck squamous cell carcinoma

### Soluble MMP2 levels were independent of T cell infiltration

MMP degrade the extra-cellular matrix and promote tumor cell invasion (15). Tissue damage may lead to a local increase in danger signals, and initiate an innate and then adaptive immune response. Thus, we hypothesized that MMP2 levels might influence T cell infiltration. Paired CD3 and CD8 T cell quantification by flow cytometry, and soluble MMP2 quantification, was available for 18 HNSCC patients. MMP2 was not significantly correlated to CD3 (r = 0.01, Spearman correlation coefficient) (Fig 2C) nor to CD8 infiltration (r = −0.13, data not shown). Conversely, CD3 and CD8 infiltration were highly correlated to CXCL9 (r = 0.78 and r = 0.79) and CXCL10 (both r = 0.66) (Fig 2C, data not shown for CD8). In the secretome analysis of the 37 OCSCC samples, MMP2 was not correlated to CXCL9 and CXCL10 (r=0.19 and r=0.09), further supporting that MMP2 levels were not associated to T cell infiltration (Fig 2D).

### RNA levels of *MMP2, CD276, CXCL10*, and *STAT1* predicted prognosis

To independently validate the prognostic value of MMP2, we measured a 30 genes panel (Table S4) by RTqPCR in a large retrospective cohort of 145 OCSCC patients treated by primary surgery. Gene panel included MMP-2, −1, −9, other immune-related genes, and a published 18-gene signature predictive of the response to anti-PD-1 immunotherapy (16). Patients’ characteristics are available in Table 1. Significant variables in univariate analysis for DSS, OS and DFS are listed in Table 2. Among the clinical variables, tumor differentiation index, stage, ENE, VE and PNI were significant for both DSS and OS, while only the latter three were significant for DFS. Among the genes, high levels of *MMP2* were associated to reduced DSS, OS and DFS. High levels of *CD276* (B7-H3) and low levels of *CXCL10* and *STAT1* were also among the 5 and 11 genes associated to reduced DSS and OS, respectively (Table 2). This validated the prognostic impact of MMP2, measured by two different methods (protein and mRNA), in a large OCSCC cohort.

**Table 1.**
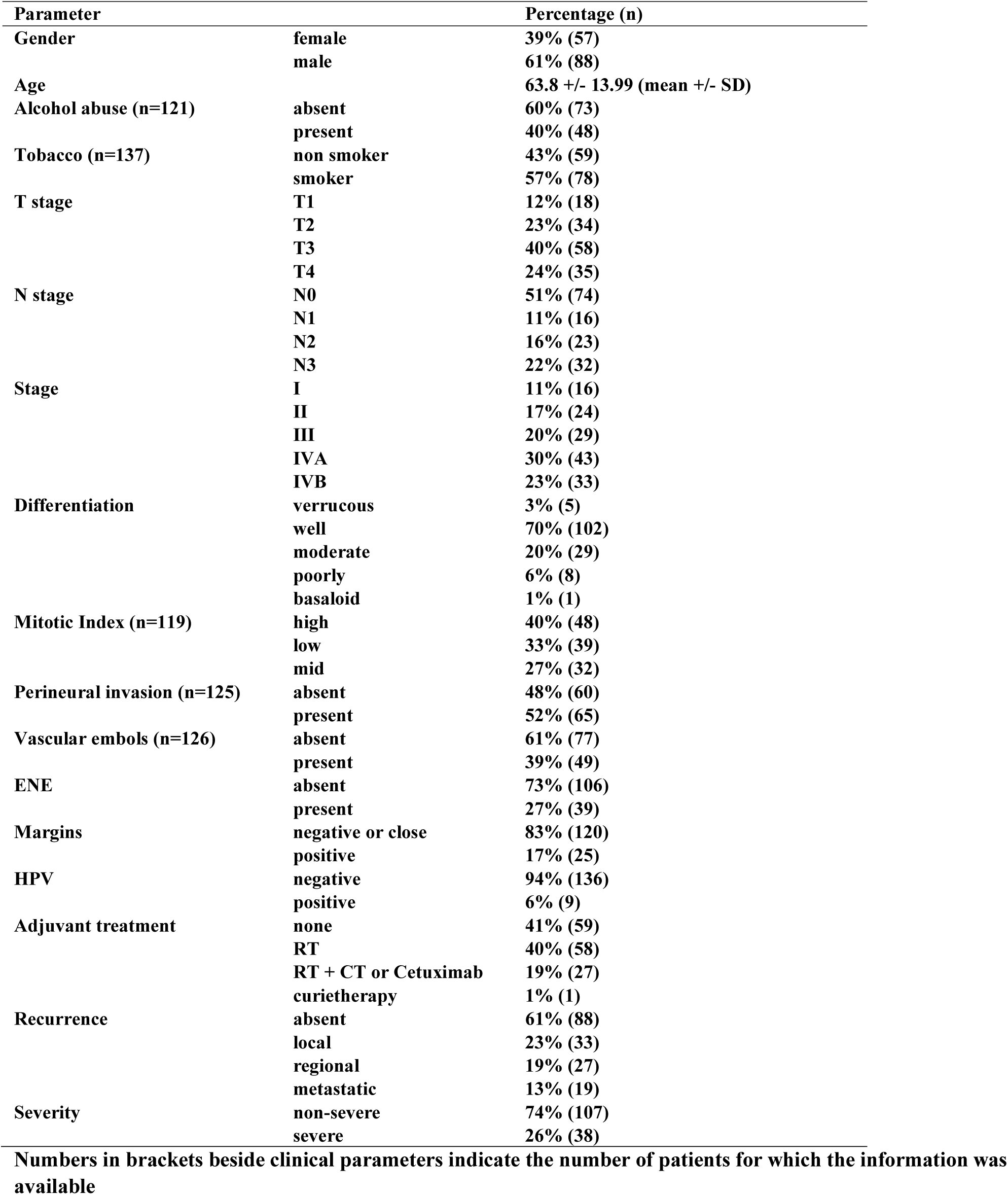
Patients characteristics of the RT-qPCR retrospective validation cohort (n=145)

**Table 2.** Prognosis value of the clinical parameters and genes measured by RTqPCR in the validation cohort (univariate analysis, Log-Rank test)

| Parameter | Mean +/- SD | Poor prognosis if | p-values per survival (Log-rank) |  |  |
| --- | --- | --- | --- | --- | --- |
|  |  |  | DSS | OS | DFS |
| Gender |  | ns | 0.8420 | 0.4387 | 0.801 |
| Age (</> 70) |  | ns | 0.9460 | 0.9785 | 0.434 |
| Alcohol |  | ns | 0.8710 | 0.1860 | 0.848 |
| Tobacco |  | ns | 0.7839 | 0.1191 | 0.670 |
| Stage |  | III or more | 0.0120 | 0.0036 | 0.053 |
| Differentiation |  | moderate or poor | 0.0350 | 0.0434 | 0.117 |
| Mitotic index |  | ns | 0.1957 | 0.7066 | 0.928 |
| Perineural invasion |  | present | < 0.0001 | < 0.0001 | 0.0046 |
| Vascular embols |  | present | 0.0004 | 0.0002 | 0.0130 |
| ENE |  | present | < 0.0001 | 0.0004 | 0.003 |
| Margins |  | ns | 0.1020 | 0.1484 | 0.193 |
| HPV |  | ns | 0.4950 | 0.4536 | 0.823 |
| MMP2 | 1.84+/-1.75 | high | 0.0009 | 0.0140 | 0.0440 |
| CD276 | 2.4+/-1.18 | high | 0.0056 | 0.0340 | 0.0870 |
| CXCL10 | 18.67+/-27.62 | low | 0.0083 | 0.0008 | 0.0820 |
| STAT1 | 3.72+/-2.35 | low | 0.0160 | 0.0007 | 0.1300 |
| MMP9 | 8.55+/-12.93 | high | 0.0190 | 0.0880 | 0.0610 |
| LAMP3 | 7.43+/-5.59 | low | 0.1500 | 0.0008 | 0.4300 |
| CXCR6 | 1.22+/-0.92 | low | 0.6200 | 0.0037 | 0.6600 |
| HLA-E | 1.12+/-0.51 | low | 0.1100 | 0.0056 | 0.0810 |
| CD274 | 3.3+/-3.25 | low | 0.2100 | 0.0070 | 0.4100 |
| IDO1 | 13.98+/-20.3 | low | 0.0650 | 0.0095 | 0.1800 |
| PSMB10 | 1.68+/-0.99 | low | 0.2000 | 0.0270 | 0.2800 |
| CCR7 | 8.41+/-10.73 | low | 0.4700 | 0.0300 | 0.5900 |
| TIGIT | 3.28+/-2.8 | ns | 0.8800 | 0.0560 | 0.7700 |
| CCL5 | 2.3+/-2.41 | ns | 0.7700 | 0.0600 | 0.8800 |
| LAG3 | 3.04+/-3.28 | ns | 0.4700 | 0.0640 | 0.7900 |
| PDCD1 | 2.19+/-2.17 | ns | 0.8500 | 0.0670 | 0.5400 |
| CXCL9 | 19.04+/-30.47 | ns | 0.7000 | 0.0680 | 0.9800 |
| HLA-DQA1 | 1.5+/-1.2 | ns | 0.5600 | 0.0850 | 0.7200 |
| IL3RA | 0.9+/-0.69 | ns | 0.6300 | 0.0990 | 0.3700 |
| CD27 | 1.88+/-2.06 | ns | 0.7700 | 0.0990 | 0.7000 |
| NKG7 | 1.83+/-2.12 | ns | 0.7900 | 0.1300 | 0.4700 |
| CD3E | 2+/-1.9 | ns | 0.8100 | 0.1400 | 0.7700 |
| pan_HLA-DRB | 1.35+/-1.04 | ns | 0.7000 | 0.1500 | 0.6300 |
| PDCD1LG2 | 2.64+/-2.24 | ns | 0.3100 | 0.2000 | 0.2200 |
| CD8A | 1.74+/-2.1 | ns | 0.6200 | 0.2800 | 0.4000 |
| ICOSLG | 0.68+/-0.35 | ns | 0.9400 | 0.4200 | 0.4600 |
| CMKLR1 | 1.13+/-0.8 | ns | 0.4200 | 0.4300 | 0.4800 |
| MMP1 | 774.76+/-1051.42 | ns | 0.3000 | 0.6300 | 0.3500 |
| FUT4 | 1.06+/-0.53 | ns | 0.1600 | 0.8600 | 0.4000 |
| CD1C | 0.36+/-0.42 | ns | 0.2300 | 0.9400 | 0.4500 |

Cells highlighted in grey contain significant values at p < 0.05

### *MMP2* RNA, ENE, PNI and stage were independent prognostic factors

To identify clinical and biological parameters significant in multivariate analysis, we performed two Cox proportional hazards models. Model 1 included all the 145 patients and all clinical and biological variables significant in univariate analysis, except PNI and VE, because of missing values in 21 patients (14%), whereas Model 2 included all significant variables, but was restricted to the 124 patients with complete data (Fig 3A, Table S5). In both models *MMP2*high was an independent prognostic factor for DSS and DFS (Model 1 DSS: p = 0.001, DFS: p = 0.006, Model 2 DSS: p = 0.034, DFS: p = 0.016). For DSS, ENE status (p = 0.006) and PNI (p = 0.020) were also significant in Model 1 and 2, respectively. For DFS, ENE status was also significant in Model 1 (p = 0.006), but *MMP2* was the only significant parameter in Model 2. For OS, *MMP2* (p = 0.015) and stage (p = 0.042) were significant in Model 1, and PNI (p = 0.01) and stage (p = 0.019) were significant in Model 2 (Fig 3A, Table S5). We defined prognostic groups using the parameters identified in multivariate analysis by the Model 1 to analyze the largest cohort of 145 patients. *MMP2*highENE+ patients had the worse DSS and DFS, as compared to *MMP2*lowENE-patients (p < 0.001), whereas *MMP2*highENE- and *MMP2*lowENE+ had an intermediate DSS and DFS (Fig 3B) (2 by 2 comparisons available in Table S6). *MMP2* status induced clinically relevant variations in survival. *MMP2*high vs *MMP2*low tumor bearing patients had a 5-year DSS of 61% versus 88% when ENE was absent, and of 36% versus 52% when ENE was present (Table 3). *MMP2*high tumors were associated to the presence of metastatic lymph node (p = 0.031), low or intermediate mitotic index (p = 0.001) and the presence of PNI (p = 0.02) (Table S7).

**Table 3.**
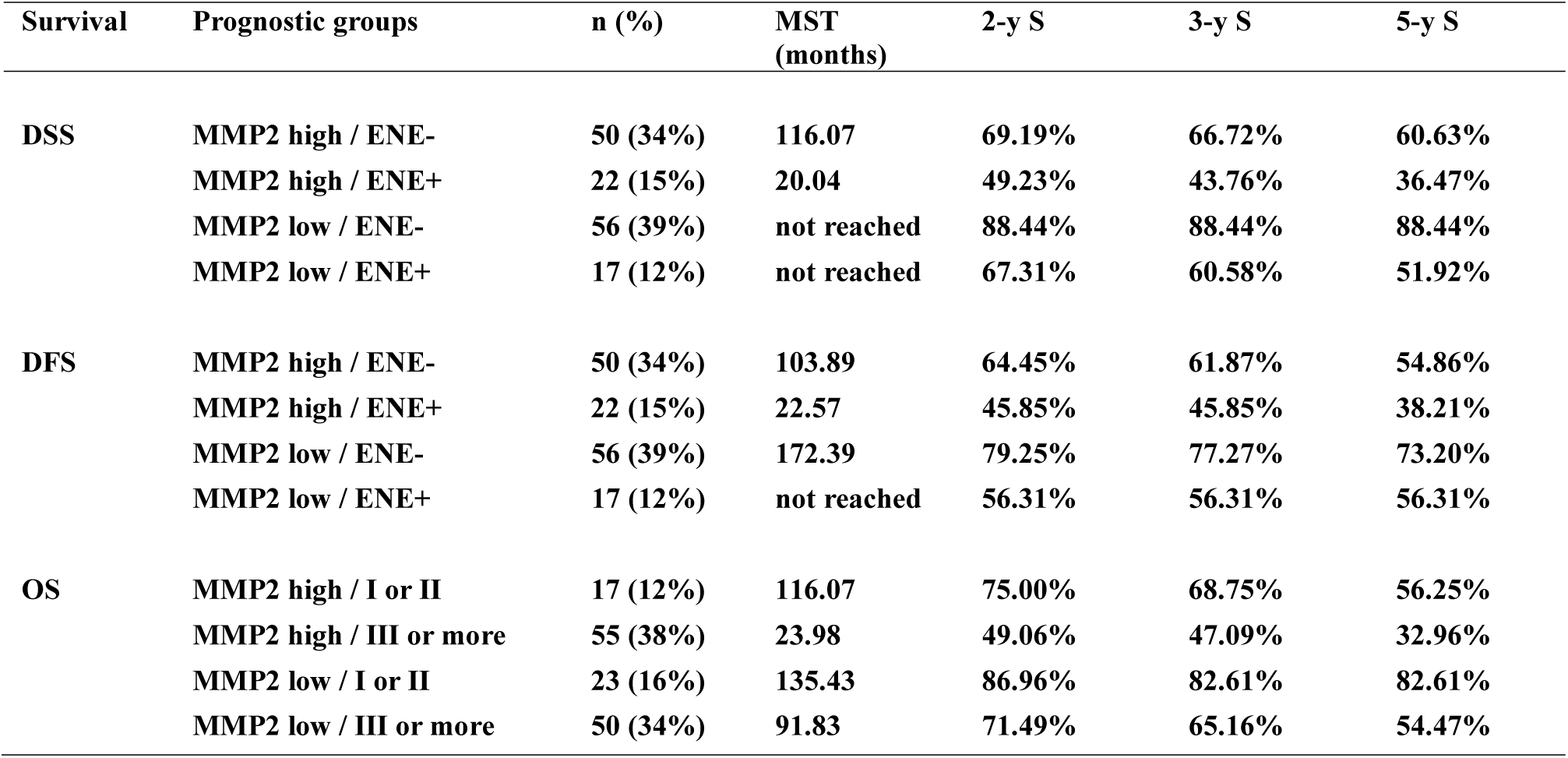
Survival durations by prognostic groups defined by the Cox Model 1

**Fig 3.**
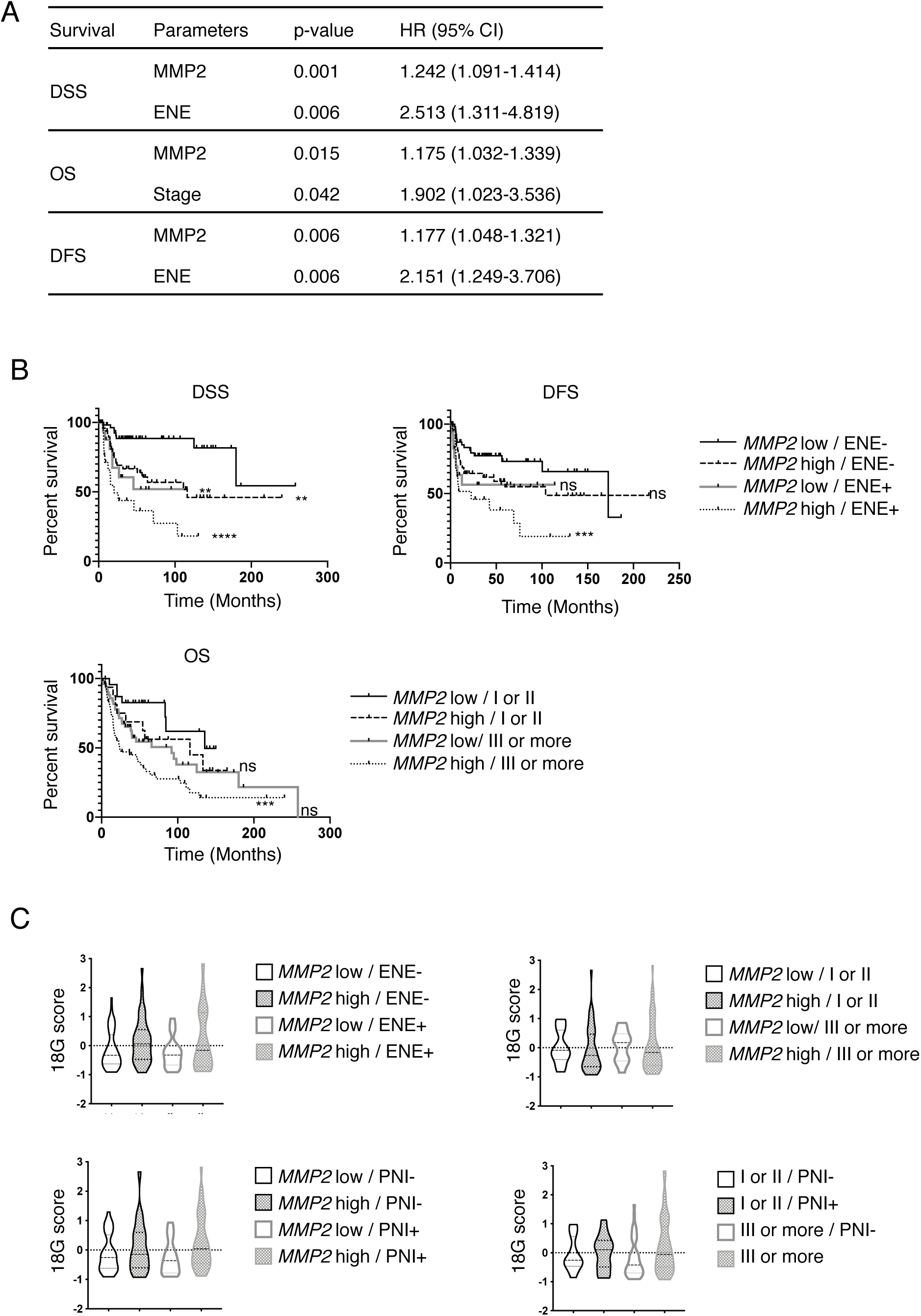
*MMP2*, ENE and stage define prognostic groups with equivalent expression of an 18-gene signature predictive of response to PD-1 blockade. A. Cox proportional hazards Model 1, including n = 145 patients, and all clinical and biological data significant at p < 0.05 in univariate analysis, excepted perineural invasion and vascular embols. B. Survivals according to the prognostic groups defined by the Cox Model 1: DSS (top left) and DFS (top right) in the 4 groups defined by *MMP2* RNA and ENE status. OS (bottom) in the 4 groups defined by MMP2 status and Stage. P-value obtained by Log-rank tests are represented by range: * < 0.05, ** < 0.01, *** < 0.001, **** < 0.0001, and relatively to the best prognosis groups that are *MMP2* low/ENE-for DSS and DFS, and *MMP2* low/Stage I or II for OS. C. Distribution of the 18-gene signature score among the prognostic groups defined by the Cox Model 1 and 2 for DFS, DSS and OS. Abbreviations. DSS: disease specific survival, DFS: disease free survival, ENE: extranodal extension, OS: overall survival.

### MMP2 may be used as a biomarker to select patients for treatment intensification

*MMP2* RNA status was an efficient prognostic biomarker as measured by ROC curves according to severity criteria, in the whole 145 patient cohort (AUC = 0.66, p = 0.003), and among the ENE negative patients (n = 106, AUC = 0.71, p = 0.003) (Fig S1). The optimal thresholds were 1.81 and 1.82, which led to high negative predictive values (NPV) of 82% and 88% respectively, but lower positive predictive values (PPV) of 41% and 36%. For 29 patients, both soluble MMP2 and *MMP2* RNA data were available, which allowed us to observe that both biomarkers were significantly correlated (Spearman r = 0.45, p = 0.016) (Fig S2), suggesting that MMP2 protein or RNA levels can be used as biomarker.

### The expression of an 18-gene signature predictive of response to PD-1 blockade was similar between the different prognostic groups

The proportions of patients expected to respond to immunotherapy may vary between the prognostic groups defined above, and have consequences on the type of treatment that could be proposed in a risk-adjusted strategy. Therefore, we measured the expression of an 18-gene signature (18G) (16) that is a predictive biomarker of response to PD-1 blockade. The 18G signature is composed of a core of 17 highly correlated genes (all Spearman correlation coefficients of the 17genes > 0.455), and CD276 (Fig S3, Fig S4). 18G score was moderately increased in *MMP2*high tumors (p = 0.019) (Fig S4, Fig S5), but was similar whatever the ENE status (p=0,671) and disease stage (p = 0.513) (Fig S5). The 18G score was similar between the prognostic groups defined by *MMP2* RNA and ENE status (p=0.119), *MMP2* RNA status and Stage (p = 0.051), *MMP2* RNA and PNI statuses (p = 0.089), and stage and PNI status (p = 0.661) (Fig 3C). This suggests that various prognostic groups may show response to anti-PD-1 therapy, with implications for the design of biomarker-driven trials in untreated resectable OCSCC patient with the goal of limiting early and severe recurrences (Fig S6).

## DISCUSSION

In this study, we identified MMP2 as an independent prognostic biomarker for severe outcomes in OCSCC patients treated by primary surgery.

First, we prospectively produced and analyzed tumor and juxtatumor secretomes, which revealed 29 deregulated soluble molecules, the majority of them being upregulated in the tumor tissue. Those molecules belonged to various biological classes such as MMPs, chemokines, interleukins, adipokines and growth factors. One may consider that all these deregulated proteins reflect mechanisms of tumor progression, and could be candidate biomarkers. However, only soluble MMP2 was associated to poor prognosis in our study. Primary tumor-derived supernatant is not a widely applied method for biomarker identification and data on OCSCC secretome are scarce (17) if we exclude cancer cell-line derived supernatants. A database for healthy body fluids proteome was created in 2008, highlighting the general interest for such approach (18). Here, we cannot exclude that tissue handling, although limited to the minimum in our protocol, may have induced or enhanced the production of some proteins, but this limitation was partially overcome by the comparison with paired juxtatumor supernatant. By the mean of an ultrafiltration catheter, interstitial fluid from a single HNSCC patient was analyzed and revealed 525 proteins by mass spectrometry, but the method was not applicable to juxtatumor tissue, which limited the potential to identify candidate biomarkers (19). Another difficulty is that tumor secretome needs to be produced prospectively using fresh tumor samples, which limits the access to large cohorts with sufficient follow-up in order to identify prognostic biomarkers. However, we could overcome these difficulties, and our study illustrates the added value of this approach in providing data with strong biological relevance. For further validation, we designed a homogenous retrospective cohort of patients with the same clinical setting of resectable OCSCC treated by primary surgery, and extracted tumor RNA from biobanked frozen samples to ensure the best quality of RNA (20). Univariate analysis confirmed the prognostic value of *MMP2* to predict DSS, OS and DFS. High levels of *CD276* and low levels of *CXCL10* and *STAT1* were also associated to reduced DSS and OS, but only *MMP2* remained significant in multivariate analysis. Several studies have proposed MMP2 as a prognostic biomarker for OCSCC, but all had important limitations, such as the absence of multivariate analysis (21), (22), (23), the inclusion of heterogeneous head and neck cancer patients with different tumor locations and treatments (24), (25), or retrospective cohorts with less than 60 patients (22), (23), (26), (27). Most of these studies quantified MMP2 by immunohistochemistry (IHC) through semi-quantitative methods. Our study provided unbiased and definite evidence for the independent prognostic role of MMP2, in a large homogeneous OCSCC cohort, within a multivariate prognostic model.

The biological basis explaining why MMP2 is associated with poor prognosis is well known. MMP2 degrades type IV collagen and promotes epithelial-mesenchymal transition and metastasis (15), (28). MMP may also skew the anti-tumor immune response by their effect on immune cells (29). MMP2 is secreted in an inactive form (pro-MMP2) and is activated by MMP1 (30) and MMP14 (31). Many cell types may produce MMP2, but fibroblasts seem to be the main source of this molecule in the tumor microenvironment (32), (33). From MMP biology, we understand that a high level of MMP is a risk factor for cancer-related events, such as recurrence and disease-related death. This explains why in our study the accuracy of MMP2 as prognostic biomarker was better for DSS than for OS, both in univariate and multivariate analysis. It is well known that HNSCC patients have a reduced cancer-independent life expectancy, which explains the differences observed between OS and DSS (34). In this line, in the TCGA data, MMP2 was co-expressed with MMP1, MMP9 and MMP14 in HNSCC, but the authors did not report the impact of any MMP on OS in HNSCC (35). The absence of DSS evaluation may explain this discrepancy. Beyond prognosis, MMP were also candidate therapeutic targets in cancer, but, so far, most molecules failed in their development because of their toxicities (36). Selective inhibitors are still in development (37), (NCT03486730), as well as other drugs that have an indirect effect on MMP (38). Clinical and histopathological parameters fail to identify around 25% of high-risk patients. Here, we propose that combining MMP2 status to those parameters would improve patients’ risk stratification. MMP2-high tumor bearing patients could be proposed for an intensified therapeutic plan, as compared to standard of care. MMP2 status may be defined pre-operatively on the initial biopsy, or post-operatively if analyzed on the resection specimen (Fig S6). Pre-operative stratification would guide neoadjuvant treatment such as immunotherapy or chemotherapy, when post-operative stratification would guide adjuvant treatment. The latter setting is particularly important for ENE negative patients who may, in some cases, not be offered any adjuvant treatment. To address the question of the best (neo)adjuvant treatment option in high risk patients, we measured the expression of an 18-gene signature predictive of response to PD-1 blockade. This signature was established on a large cohort of patients treated by pembrolizumab for head and neck cancers (n=107), melanoma (n=89) and other cancers (n=119) (16). The fact that this signature was established by merging the data from 22 different types of cancers and limited to advanced and recurrent cancers might not reflect the clinical setting of the present study. However, *PDL1* and interferon gamma response genes (*STAT1, CXCL9, IDO1, HLADR, HLADQ*) were part of this 18-gene signature and were identified as predictive of response to neoadjuvant pembrolizumab in a window-of-opportunity trial including untreated head and neck cancer patients (13). Therefore, this 18G signature may be used to estimate expected response rates to PD-1 blockade of untreated OCSCC. There was no difference in expression of the 18G score among the different prognostic groups defined by our multivariate analysis for DSS, DFS and OS. In this line, using soluble CXCL9 and CXCL10 as surrogates for tumor T cell infiltration, or direct measures of frequencies of tumor-infiltrating T cells by flow cytometry, we observed that soluble MMP2 levels were not associated to T cell infiltration. Similar results were previously described for MMP2 measured by IHC in endometrial cancer (39). From these results, we may estimate that the proportion of patients expected to respond to PD-1 blockade should be similar in the different prognostic groups, leaving immunotherapy as a valid treatment option. Patient stratification in future OCSCC trials and clinical practice would definitely benefit from robust biomarkers used in combination with clinical variables, such as our MMP2 / ENE scoring, and with predictive biomarkers for final treatment decision-making.

## MATERIALS AND METHODS

### Patients and cohorts

Tumor and juxtatumor samples were obtained from operative specimens from previously untreated head and neck cancer patients. Patients with previous head and neck radiotherapy or chemotherapy were excluded. Juxta-tumor samples were taken on the specimens’ margins, at least 1cm away from the tumor. Three cohorts of patients treated in our anti-cancer center were included in this study. All analysis on secretome presented in Fig.1 were done on a 37 patient cohort including OCSCC patients only, with the exception of the 3 graphs of Fig1D that show the correlation of CD3 infiltration with soluble MMP2, CXCL9 and CXL10, that was done in a 18 patients HNSCC cohort. This 18 patient cohort had paired secretome and flow cytometry data available and included the following tumor locations: 8 oral cavity, 6 oropharynx, 3 larynx, 1 hypopharynx. The third cohort included 145 OCSCC patients and was used to analyze gene expression by RTqPCR and prognosis. Twenty-nine patients were in common between the n=37 and n=145 cohorts and served for the RNA versus soluble protein correlation. Patients were treated between March 2010 and October 2016, for the 37 patients cohort, between January and July 2017 for the 18 patients cohort, and between February 1991 and November 2016 for the 145 patients cohort. The clinical parameters analyzed were all binarized as follows: gender (male/female), HPV status (positive by PCR/negative), Differentiation (well differentiated or verrucous or basaloid / moderate or poor), Mitotic index (high if ≥10mitoses/field at X400, otherwise low), Perineural invasion (absent/present), Vascular embols (absent/present), Alcohol (positive if ≥30g/day), Tobacco (smoker active or former ≥2PY/non-smoker or former smoker < 2PY), Stage (I or II / III or more) using the pTNM 8^th^ edition AJCC (40), Extranodal extension (absent/present), Margins (negative or close / positive), Age (more or less than 70). For outcomes analysis, we used 3 survivals: disease free survival, in which the censoring event was the first occurrence of recurrence, disease specific survival, in which the censoring event was the occurrence of death caused by the evolution of the cancer (to the exclusion of treatment related toxicities and post-operative complications), and overall survival. We also used a binary criteria of severity defined as present in cases of DSS < 36 months and /or a DFS < 12 months without subsequent remission (unsuccessful salvage procedures and/or permanent palliative treatment); we considered that these criteria define the population with the most urgent need for prognosis biomarkers (41). This study was done in compliance with the principles of Good Clinical Practice and the Declaration of Helsinki. All patients signed a consent form mentioning that their operative specimens might be used for scientific purposes, and 12 of the 18 patients cohort were also included in the clinical trial NCT03017573.

### Tumor and juxta-tumor secretome analysis

Fresh tumor and juxta-tumor were cut into fragments of 17.5 +/-2.5mg. Each fragment was placed in a 48-well flat bottom plate in 250µl of RPMI 1640 Medium Glutamax (Life Technologies) enriched with 10% Fetal Calf Serum (Hyclone), 100 U/ml Penicillin/Streptomycin (Gibco), 1% MEM Non-Essential Amino Acids (Gibco), and 1% pyruvate (Gibco), and incubated at 37°C with 5%CO2. After 24 hours, supernatants were filtered through a 0,22µm Millex-GP filter (SLGP033RS, Merck), diluted ½ in the same enriched RPMI Medium and stored at −80°C until the secretome analysis. The 49 analytes measured are listed in Table S2. Analytes concentrations were obtained using Milliplex Map kits used as recommended: Human MMP magnetic Bead panel 2, Human cytokine/chemokine Magnetic Bead panels I, II, III, and Human Adipocyte Magnetic Bead Panel (Millipore), a Bio-Plex 200 plate reader and the Bio-Plex Manager 6.1 software (Bio-Rad Laboratories). All analytes were measured as stored, but MMP1 and MMP9 were also measured after 1/25th dilution for the 18 HNSCC patients with paired flow cytometry data.

### Analysis of CD3 and CD8 infiltration by Flow Cytometry

Details are available at (42). Briefly, single-cell suspensions were obtained from enzymatically digested tumor samples, then filtered, washed, counted and stained for 15 minutes with DAPI (Miltenyi Biotec) to exclude dead cells, CD3 (Alexa700, clone UCHT1, from BD, #557943) and CD8b (PC5, clone 2ST8.5H7, from Beckman Coulter, #6607109) antibodies, among other antibodies (data not used in the present paper), before phenotyping by flow cytometry (BD LSRFortessa Analyzer).

### Gene expression analysis by Real-Time RT-PCR

#### Samples and RNA Extraction

Tumor and juxtatumor samples were snap-frozen in liquid nitrogen upon surgical removal after pathologist’s review and were stored in the corresponding our biological resources center. Samples were sectioned using Tissue-Tek optimal cutting temperature (O.C.T) compound to estimate the percentage of tumor cells and to remove non-malignant tissue by macrodissection if necessary. Median percentage of tumor cells was 80% (range 40-95). RNA extraction was performed on the same sample, using the miRNeasy miniKit (Qiagen) according to the manufacturer’s protocol. RNA was quantified using Nanodrop spectrophotometer ND-1000 and the integrity and purity were assessed by the Agilent 2100 Bioanalyzer and RNA 6000 Nano Labchip Kit (Agilent Biotechnologies, Palo Alto, CA, USA).

Total RNA was extracted from 145 OCSSC and 31 juxtatumor frozen samples from HNSCC bearing patients by using the acid-phenol guanidium method. RNA samples quality was assessed by electrophoresis through agarose gels and staining with ethidium bromide, and the 18S and 28S RNA bands were visualized under UV light.

#### cDNA Synthesis

RNA was reverse transcribed in a final volume of 20 μl containing 1X RT buffer, 0.01M DTT, 0.5mM each dNTP, 0.15µg/µL random primers, 100U SuperScript™ II Reverse Transcriptase (Life Technologies, Carlsbad, Californie), 20U RNasin® Ribonuclease Inhibitor (Promega, Madison, Wisconsin) and 1 μg of total RNA. The samples were incubated during 10min at 25°C 30min at 42°C, and reverse transcriptase was inactivated by heating 5min at 99°C and cooling 5min at 5°C.

#### PCR Amplification and quantification

All of the PCR reactions were performed using an ABI Prism 7900HT Sequence Detection system (Thermo Fisher Scientific, Waltham, Massachusetts). PCR was performed using the *Power* SYBR™ Green PCR Master Mix (Life Technologies, Carlsbad, Californie). The thermal cycling conditions comprised an initial denaturation step of 10min at 95°C followed by 50 cycles at 95°C for 15 s and 65°C for 1 min. Cycle Threshold (Ct value) was defined by the cycle number at which the increase in the fluorescence signal associated with exponential growth of PCR products started to be detected, using Applied Biosystems analysis software according to the manufacturer’s manuals. For quality controls, we quantified the housekeeping gene TBP (Genbank accession NM_003194). Primers for TBP and the 30 target genes were designed with the assistance of Oligo 6.0 computer program (National Biosciences, Plymouth, MN). dbEST and nr databases were used to confirm the total gene specificity of the nucleotide sequences chosen as primers and the absence of single nucleotide polymorphisms. The primer pairs selected were unique relative to the sequences of closely related family member genes and the corresponding retropseudogenes. One of the two primers was placed at the junction between two exons or on two different exons to avoid genomic DNA contaminating. Specificity of PCR amplicons was verified by agarose gel electrophoresis. The oligonucleotide primers sequences used are shown in Table S8.

#### Data processing

TBP was used for each sample normalization. ΔCt value was equal to mean Ct value of the target gene minus mean Ct value of TBP. The N-fold differences per sample in target gene expression relative to TBP was equal to 2^ΔCt^. For each gene, 2^ΔCt^ values of the 31 juxtatumor samples were multiplied by a factor named “k” so that their median was equal to 1. The final values for tumor samples were equal to k2^ΔCt^. The 30 genes of this study are listed in Table S5. To obtain a score for the 18 genes signature, we standardized each gene separately, and used those values in the formula: 18G score = (*CCR7+ HLADRB + CCL5 + CD27 - CD276 + CMKLR1 + CXCL9 + CXCR6 + HLA-DQA1 + HLA-E + IDO1 + LAG3 + NKG7 + PDCD1LG2 + PSMB10 + STAT1 + TIGIT*)/18.

### Statistical Analysis

Descriptive and statistical analyses were performed using GraphPad Prism V8, Xlstat (Addinsoft), and Qlucore softwares. Paired tumor and juxtatumor secretome comparison was done by Wilcoxon test. Univariate unpaired non-parametric comparisons used Mann-Whitney tests and Kruskal-Wallis test for multigroup comparisons. All correlations used Spearman method. Optimal threshold for ROC curves was defined as the value maximizing the sum of sensitivity and specificity. Univariate survival analysis was performed on clinical parameters and biological parameters (soluble molecules or 30 genes measured by RT-PCR) categorized as high or low by cut-off at median, or at optimal threshold when specified. Log-rank tests were used for univariate analysis. For the 145 patient validation cohort, significant variables at the threshold of p < 0.05 were selected for the Cox proportional hazard models for multivariate analysis. Model 1 included 145 patients and all clinical and biological parameters significant in univariate analysis, but PNI and VE, because of missing values, whereas Model 2 included all significant parameters, but was restricted to the 124 patients with complete data. The heatmap representing the 18-gene signature in Fig3A was performed with Qlucore software.

## SUPPORTING INFORMATION

“This work was supported by the Institut National de la Santé et de la Recherche Médicale under Grants BIO2012-02, BIO2014-08, and HTE201606; Agence Nationale de la Recherche under Grants ANR-10-IDEX-0001-02 PSL*, ANR-11-LABX-0043 CIC IGR-Curie 1428; Institut National du Cancer under Grant Cancéropole INCA PhD grant to CH, and INCA PLBio Grant (INCA 2016-1-PL BIO-02-ICR-1); Ligue nationale contre le cancer (labellisation EL2016.LNCC/VaS); and Institut Curie, in particular the PIC TME.

## ACKNOWLEDGMENTS

The authors wish to thank the INSERM U932, and the Institut Curie Flow-Cytometry facility, in particular Olivier Lantz, Zofia Maciorowsky, Annick Viguier, and Sophie Grondin for their technical help and expertise.

## SUPPLEMENTARY FIGURE LEGENDS

Fig S1. ROC curve of *MMP2* RNA for severity criteria in the cohort of 145 patients (left) and among the 106 patients without ENE (right).

Fig S2. Correlation between soluble MMP2 and *MMP2* RNA (Spearman correlation coefficient).

Fig S3. Heatmap representing the expression of the 18 genes of the signature ordered by the 18-gene signature score from low values (left) to high values (right).

Fig S4. Correlation matrix of *MMP2* RNA and the genes of the 18-gene signature (Spearman correlation coefficient).

Fig S5. Distribution of the 18-gene signature score among *MMP2* RNA high and low tumors (left), absence or presence of ENE (center) and disease stage (right).

Fig S6. Flow-chart representing a proposal of an MMP2-driven clinical trial

## REFERENCES

1. NCCN. https://www.nccn.org/professionals/physician_gls/default.aspx.

2. Gañán L, López M, García J, Esteller E, Quer M, León X. Management of recurrent head and neck cancer: variables related to salvage surgery. Eur Arch Oto-Rhino-Laryngol Off J Eur Fed Oto-Rhino-Laryngol Soc EUFOS Affil Ger Soc Oto-Rhino-Laryngol - Head Neck Surg. 2016 Dec;273(12):4417–24.

3. Argiris A, Karamouzis MV, Raben D, Ferris RL. Head and neck cancer. Lancet Lond Engl. 2008 May 17;371(9625):1695–709.

4. Tam S, Araslanova R, Low T-HH, Warner A, Yoo J, Fung K, et al. Estimating Survival After Salvage Surgery for Recurrent Oral Cavity Cancer. JAMA Otolaryngol -- Head Neck Surg. 2017 01;143(7):685–90.

5. Ord RA, Kolokythas A, Reynolds MA. Surgical salvage for local and regional recurrence in oral cancer. J Oral Maxillofac Surg Off J Am Assoc Oral Maxillofac Surg. 2006 Sep;64(9):1409–14.

6. Sacco AG, Cohen EE. Current Treatment Options for Recurrent or Metastatic Head and Neck Squamous Cell Carcinoma. J Clin Oncol Off J Am Soc Clin Oncol. 2015 Oct 10;33(29):3305–13.

7. Liao C-T, Chang JT-C, Wang H-M, Ng S-H, Hsueh C, Lee L-Y, et al. Salvage therapy in relapsed squamous cell carcinoma of the oral cavity: how and when? Cancer. 2008 Jan 1;112(1):94–103.

8. Janot F, de Raucourt D, Benhamou E, Ferron C, Dolivet G, Bensadoun R-J, et al. Randomized trial of postoperative reirradiation combined with chemotherapy after salvage surgery compared with salvage surgery alone in head and neck carcinoma. J Clin Oncol Off J Am Soc Clin Oncol. 2008 Dec 1;26(34):5518–23.

9. Zorat PL, Paccagnella A, Cavaniglia G, Loreggian L, Gava A, Mione CA, et al. Randomized phase III trial of neoadjuvant chemotherapy in head and neck cancer: 10-year follow-up. J Natl Cancer Inst. 2004 Nov 17;96(22):1714–7.

10. Bossi P, Lo Vullo S, Guzzo M, Mariani L, Granata R, Orlandi E, et al. Preoperative chemotherapy in advanced resectable OCSCC: long-term results of a randomized phase III trial. Ann Oncol Off J Eur Soc Med Oncol. 2014 Feb;25(2):462–6.

11. Uppaluri R, Zolkind P, Lin T, Nussenbaum B, Jackson RS, Rich J. Neoadjuvant pembrolizumab in surgically resectable, locally advanced HPV negative head and neck squamous cell carcinoma (HNSCC). In: J Clin Oncol 35, 2017 (suppl; abstr 6012).

12. Ferris RL, Gonçalves A, Baxi S, Martens A, Gauthier H, Langenberg M et al. An Open-label, Multicohort, Phase 1/2 Study in Patients With Virus-Associated Cancers (CheckMate 358): Safety and Efficacy of Neoadjuvant Nivolumab in Squamous Cell Carcinoma of the Head and Neck. In: Poster LBA46, ESMO 2017.

13. Wise-Draper TM, Matthew O. Old, Francis P. Worden, Paul E. O’Brien, Ezra E.W. Cohen, Neal Dunlap, Michelle Lynn Mierzwa, Keith Casper, Sarah Palackdharry, Benjamin Hinrichs, Alfredo Molinolo, Vinita Takiar, Jonathan Mark, Alice Tang, Muhammad Kashif Riaz, John Charles Morris, Nooshin Hashemi Sadraei, Changchun Xie, Maura L. Gillison. Phase II multi-site investigation of neoadjuvant pembrolizumab and adjuvant concurrent radiation and pembrolizumab with or without cisplatin in resected head and neck squamous cell carcinoma. In: ASCO 2018, abstract 6017.

14. Rivera C, Oliveira AK, Costa RAP, De Rossi T, Paes Leme AF. Prognostic biomarkers in oral squamous cell carcinoma: A systematic review. Oral Oncol. 2017;72:38–47.

15. Kessenbrock K, Plaks V, Werb Z. Matrix metalloproteinases: regulators of the tumor microenvironment. Cell. 2010 Apr 2;141(1):52–67.

16. Cristescu R, Mogg R, Ayers M, Albright A, Murphy E, Yearley J, et al. Pan-tumor genomic biomarkers for PD-1 checkpoint blockade-based immunotherapy. Science. 2018 12;362(6411).

17. Gromov P, Gromova I, Olsen CJ, Timmermans-Wielenga V, Talman M-L, Serizawa RR, et al. Tumor interstitial fluid - a treasure trove of cancer biomarkers. Biochim Biophys Acta. 2013 Nov;1834(11):2259–70.

18. Li S-J, Peng M, Li H, Liu B-S, Wang C, Wu J-R, et al. Sys-BodyFluid: a systematical database for human body fluid proteome research. Nucleic Acids Res. 2009 Jan;37(Database issue):D907–912.

19. Stone MD, Odland RM, McGowan T, Onsongo G, Tang C, Rhodus NL, et al. Novel In Situ Collection of Tumor Interstitial Fluid from a Head and Neck Squamous Carcinoma Reveals a Unique Proteome with Diagnostic Potential. Clin Proteomics. 2010 Sep;6(3):75–82.

20. Li J, Fu C, Speed TP, Wang W, Symmans WF. Accurate RNA Sequencing From Formalin-Fixed Cancer Tissue To Represent High-Quality Transcriptome From Frozen Tissue. JCO Precis Oncol. 2018;2018.

21. Chen C-H, Chien C-Y, Huang C-C, Hwang C-F, Chuang H-C, Fang F-M, et al. Expression of FLJ10540 is correlated with aggressiveness of oral cavity squamous cell carcinoma by stimulating cell migration and invasion through increased FOXM1 and MMP-2 activity. Oncogene. 2009 Jul 30;28(30):2723–37.

22. Fan H-X, Li H-X, Chen D, Gao Z-X, Zheng J-H. Changes in the expression of MMP2, MMP9, and ColIV in stromal cells in oral squamous tongue cell carcinoma: relationships and prognostic implications. J Exp Clin Cancer Res CR. 2012 Oct 29;31:90.

23. Aparna M, Rao L, Kunhikatta V, Radhakrishnan R. The role of MMP-2 and MMP-9 as prognostic markers in the early stages of tongue squamous cell carcinoma. J Oral Pathol Med Off Publ Int Assoc Oral Pathol Am Acad Oral Pathol. 2015 May;44(5):345–52.

24. Ruokolainen H, Pääkkö P, Turpeenniemi-Hujanen T. Tissue and circulating immunoreactive protein for MMP-2 and TIMP-2 in head and neck squamous cell carcinoma--tissue immunoreactivity predicts aggressive clinical course. Mod Pathol Off J U S Can Acad Pathol Inc. 2006 Feb;19(2):208–17.

25. Gunawardena I, Arendse M, Jameson MB, Plank LD, Gregor RT. Prognostic molecular markers in head and neck squamous cell carcinoma in a New Zealand population: matrix metalloproteinase-2 and sialyl Lewis x antigen. ANZ J Surg. 2015 Nov;85(11):843–8.

26. Gontarz M, Wyszyńska-Pawelec G, Zapała J, Czopek J, Lazar A, Tomaszewska R. Immunohistochemical predictors in squamous cell carcinoma of the tongue and floor of the mouth. Head Neck. 2016;38 Suppl 1:E747–753.

27. Katayama A, Bandoh N, Kishibe K, Takahara M, Ogino T, Nonaka S, et al. Expressions of matrix metalloproteinases in early-stage oral squamous cell carcinoma as predictive indicators for tumor metastases and prognosis. Clin Cancer Res Off J Am Assoc Cancer Res. 2004 Jan 15;10(2):634–40.

28. Yokoyama K, Kamata N, Fujimoto R, Tsutsumi S, Tomonari M, Taki M, et al. Increased invasion and matrix metalloproteinase-2 expression by Snail-induced mesenchymal transition in squamous cell carcinomas. Int J Oncol. 2003 Apr;22(4):891–8.

29. Godefroy E, Manches O, Dréno B, Hochman T, Rolnitzky L, Labarrière N, et al. Matrix metalloproteinase-2 conditions human dendritic cells to prime inflammatory T(H)2 cells via an IL- 12- and OX40L-dependent pathway. Cancer Cell. 2011 Mar 8;19(3):333–46.

30. Sato H, Takino T, Okada Y, Cao J, Shinagawa A, Yamamoto E, et al. A matrix metalloproteinase expressed on the surface of invasive tumour cells. Nature. 1994 Jul 7;370(6484):61–5.

31. Jobin PG, Butler GS, Overall CM. New intracellular activities of matrix metalloproteinases shine in the moonlight. Biochim Biophys Acta Mol Cell Res. 2017 Nov;1864(11 Pt A):2043–55.

32. Puram SV, Tirosh I, Parikh AS, Patel AP, Yizhak K, Gillespie S, et al. Single-Cell Transcriptomic Analysis of Primary and Metastatic Tumor Ecosystems in Head and Neck Cancer. Cell. 2017 Dec 14;171(7):1611–1624.e24.

33. Zhang Z, Liu R, Jin R, Fan Y, Li T, Shuai Y, et al. Integrating Clinical and Genetic Analysis of Perineural Invasion in Head and Neck Squamous Cell Carcinoma. Front Oncol. 2019;9:434.

34. Massa ST, Cass LM, Osazuwa-Peters N, Christopher KM, Walker RJ, Varvares MA. Decreased cancer-independent life expectancy in the head and neck cancer population. Head Neck. 2017;39(9):1845–53.

35. Gobin E, Bagwell K, Wagner J, Mysona D, Sandirasegarane S, Smith N, et al. A pan-cancer perspective of matrix metalloproteases (MMP) gene expression profile and their diagnostic/prognostic potential. BMC Cancer. 2019 Jun 14;19(1):581.

36. Zhong Y, Lu Y-T, Sun Y, Shi Z-H, Li N-G, Tang Y-P, et al. Recent opportunities in matrix metalloproteinase inhibitor drug design for cancer. Expert Opin Drug Discov. 2018;13(1):75–87.

37. Scannevin RH, Alexander R, Haarlander TM, Burke SL, Singer M, Huo C, et al. Discovery of a highly selective chemical inhibitor of matrix metalloproteinase-9 (MMP-9) that allosterically inhibits zymogen activation. J Biol Chem. 2017 27;292(43):17963–74.

38. Van Tubergen EA, Banerjee R, Liu M, Vander Broek R, Light E, Kuo S, et al. Inactivation or loss of TTP promotes invasion in head and neck cancer via transcript stabilization and secretion of MMP9, MMP2, and IL-6. Clin Cancer Res Off J Am Assoc Cancer Res. 2013 Mar 1;19(5):1169–79.

39. Jedryka M, Chrobak A, Chelmonska-Soyta A, Gawron D, Halbersztadt A, Wojnar A, et al. Matrix metalloproteinase (MMP)-2 and MMP-9 expression in tumor infiltrating CD3 lymphocytes from women with endometrial cancer. Int J Gynecol Cancer Off J Int Gynecol Cancer Soc. 2012 Oct;22(8):1303–9.

40. Amin, M.B., Edge, S., Greene, F., Byrd, D.R., Brookland, R.K., Washington, M.K., Gershenwald, J.E., Compton, C.C., Hess, K.R., Sullivan, D.C., Jessup, J.M., Brierley, J.D., Gaspar, L.E., Schilsky, R.L., Balch, C.M., Winchester, D.P., Asare, E.A., Madera, M., Gress, D.M., Meyer, L.R. AJCC cancer staging manual. 8th ed. New York: Springer; 2017.

41. Agra IMG, Carvalho AL, Ulbrich FS, de Campos OD, Martins EP, Magrin J, et al. Prognostic factors in salvage surgery for recurrent oral and oropharyngeal cancer. Head Neck. 2006 Feb;28(2):107–13.

42. Hoffmann C, Noel F, Grandclaudon M, Michea P, Surun A, Faucheux L, et al. PDL1 and ICOSL discriminate human secretory and helper dendritic cells. bioRxiv [Internet]. 2019 Aug 1 [cited 2019 Aug 2]; Available from: http://biorxiv.org/lookup/doi/10.1101/721563

43. Daneshmandi S, Pourfathollah AA, Forouzandeh-Moghaddam M. Enhanced CD40 and ICOSL expression on dendritic cells surface improve anti-tumor immune responses; effectiveness of mRNA/chitosan nanoparticles. Immunopharmacol Immunotoxicol. 2018 Sep 28;1–12.

