## Supplementary Figures for "MMP2 As An Independent Prognostic Stratifier In Oral Cavity Cancers"

### Figure S1

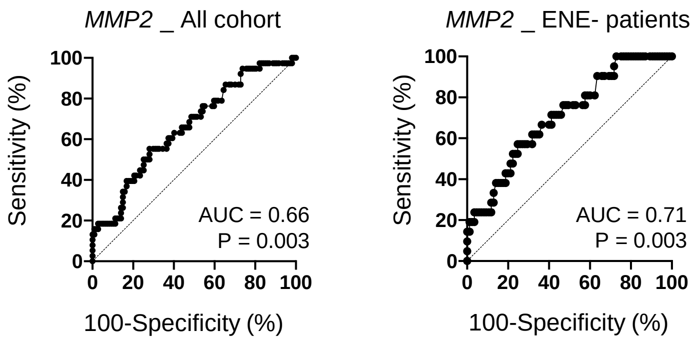
MMP2 _ All cohort
Sensitivity (%)
100-Specificity (%)
0
20
40
60
80
100
AUC = 0.66
P = 0.003
MMP2 _ ENE- patients
Sensitivity (%)
100-Specificity (%)
AUC = 0.71
P = 0.003

### Figure S2

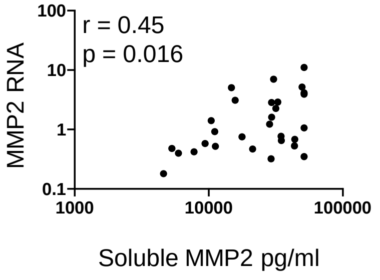

r = 0.45
p = 0.016
MMP2 RNA
100
10
1
0.1
1000
10000
100000
Soluble MMP2 pg/ml

### Figure S3

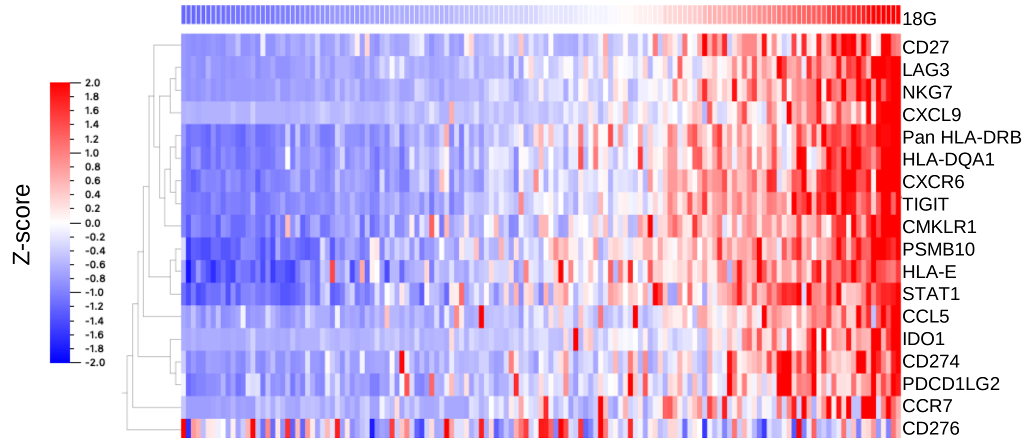
Z-score
2.0
1.8
1.6
1.4
1.2
1.0
0.8
0.6
0.4
0.2
0.0
-0.2
-0.4
-0.6
-0.8
-1.0
-1.2
-1.4
-1.6
-1.8
-2.0
18G
CD27
LAG3
NKG7
CXCL9
Pan HLA-DRB
HLA-DQA1
CXCR6
TIGIT
CMKLR1
PSMB10
HLA-E
STAT1
CCL5
IDO1
CD274
PDCD1LG2
CCR7
CD276

### Figure S4

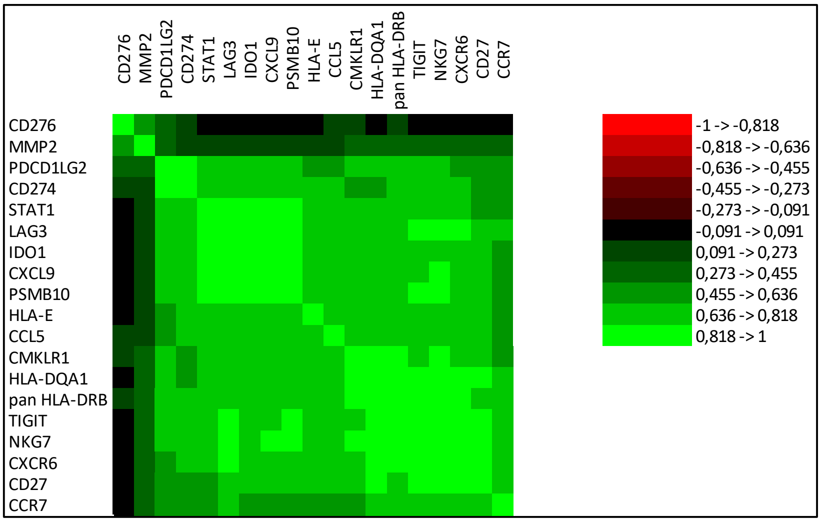
CD276
MMP2
PDCD1LG2
CD274
STAT1
LAG3
IDO1
CXCL9
PSMB10
HLA-E
CCL5
CMKLR1
HLA-DQA1
pan HLA-DRB
TIGIT
NKG7
CXCR6
CD27
CCR7
-1 -> -0,818
-0,818 -> -0,636
-0,636 -> -0,455
-0,455 -> -0,273
-0,273 -> -0,091
-0,091 -> 0,091
0,091 -> 0,273
0,273 -> 0,455
0,455 -> 0,636
0,636 -> 0,818
0,818 -> 1

### Figure S5

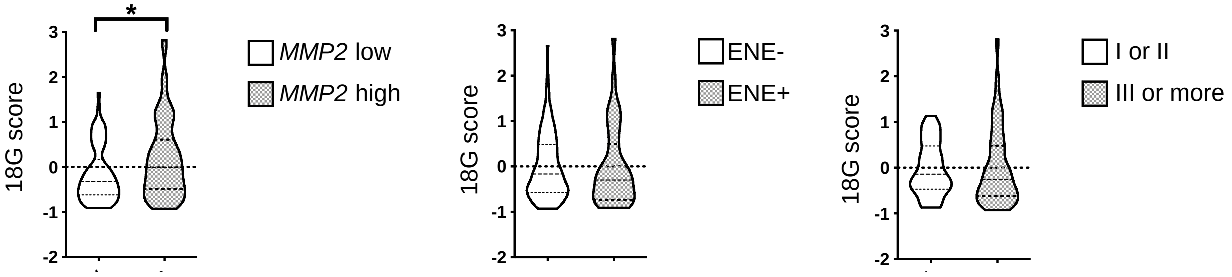

### Figure S6

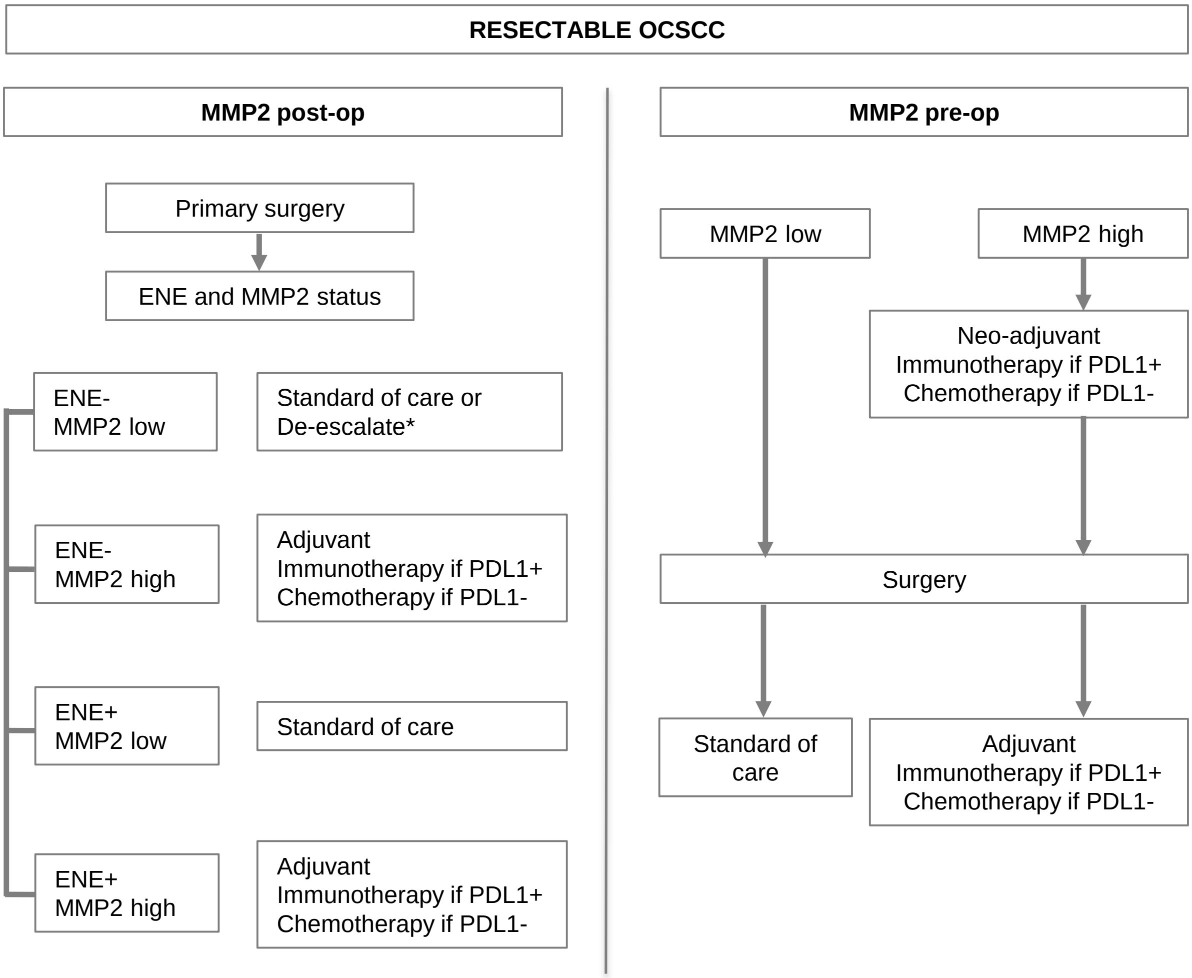


* Depending on other parameters, such as PNI, VE, N status, consider de-escalating adjuvant chemoradiotherapy to radiotherapy, or radiotherapy to surveillance
