## Supplementary Tables for "MMP2 As An Independent Prognostic Stratifier In Oral Cavity Cancers"

Table S1 - Patients characteristics of the secretome prospective discovery cohort (n = 37)

| Parameter | | Percentage (n) |
|---|---|---|
| Gender | female | 32% (12) |
| | male | 68% (25) |
| Age | | 68.31 +/- 12.81 (mean +/- SD) |
| Alcohol abuse (n=27) | absent | 67% (18) |
| | present | 33% (9) |
| Tobacco (n=34) | non smoker | 50% (17) |
| | smoker | 50% (17) |
| T stage | T1 | 14% (5) |
| | T2 | 22% (8) |
| | T3 | 32% (12) |
| | T4 | 32% (12) |
| N stage | N0 | 59% (22) |
| | N1 | 8% (3) |
| | N2 | 14% (5) |
| | N3 | 19% (7) |
| Stage | I | 14% (5) |
| | II | 11% (4) |
| | III | 19% (7) |
| | IVA | 38% (14) |
| | IVB | 19% (7) |
| Differentiation | well | 78% (29) |
| | moderate | 22% (8) |
| | poorly | 0% (0) |
| Mitotic Index (n=36) | high | 53% (19) |
| | low | 25% (9) |
| | mid | 31% (11) |
| Perineural invasion (n=36) | absent | 47% (17) |
| | present | 53% (19) |
| Vascular embols | absent | 59% (22) |
| | present | 41% (15) |
| ENE | absent | 76% (28) |
| | present | 24% (9) |
| Margins | negative or close | 86% (32) |
| | positive | 14% (5) |
| HPV (n=21) | negative | 90% (19) |
| | positive | 10% (2) |
| Adjuvant treatment | none | 30% (11) |
| | RT | 54% (20) |
| | RT + CT or Cetuximab | 16% (6) |
| Recurrence | absent | 73% (27) |
| | present | 27% (10) |
| Severity | non-severe | 76% (28) |
| | severe | 24% (9) |

Numbers in brackets beside clinical parameters indicate the number of patients for which the information was available

Table S2 - Comparison of the analytes of the soluble microenvironment of 37 paired OCSCC and juxtatumor samples (Wilcoxon)

| Analyte | Tumor Median (min-max) | Juxtatumor Median (min-max) | Higher in | p-value |
|---|---|---|---|---|
| CXCL9 | 35380 (61-52000) | 2934 (31-52000) | Tumor | <0.0001 |
| GM-CSF | 1093 (0-10800) | 105 (0-3386) | Tumor | <0.0001 |
| IL-15 | 5 (0-17) | 1 (0-8) | Tumor | <0.0001 |
| MMP-2 | 28457 (4155-51500) | 5414 (0-51500) | Tumor | <0.0001 |
| MMP-9 | 10500 (783-10500) | 2522 (159-10500) | Tumor | <0.0001 |
| PAI-1 | 19392 (1513-34000) | 4579 (61-34000) | Tumor | <0.0001 |
| Resistin | 10460 (109-24500) | 1263 (27-24500) | Tumor | <0.0001 |
| SCF | 22 (0-242) | 9 (0-42) | Tumor | <0.0001 |
| TNFa | 83 (1-2402) | 37 (0-330) | Tumor | <0.0001 |
| MCP-1 | 1103 (163-19500) | 10669 (0-19500) | Juxtatumor | <0.0001 |
| IL-1b | 843 (1-5996) | 163 (0-3221) | Tumor | 0.0001 |
| IL-12(p40) | 0 (0-24) | 0 (0-8) | Tumor | 0.0002 |
| IL-16 | 143 (18-2085) | 35 (0-632) | Tumor | 0.0003 |
| TARC | 4 (0-87) | 0 (0-15) | Tumor | 0.0003 |
| TRAIL | 17 (0-238) | 6 (0-136) | Tumor | 0.0003 |
| VEGF | 72 (0-2399) | 39 (0-228) | Tumor | 0.0006 |
| MMP-1 | 21000 (7281-21000) | 21000 (28-21000) | Tumor | 0.0024 |
| IL-12(p70) | 1 (0-14) | 0 (0-2) | Tumor | 0.0029 |
| MCP-3 | 0 (0-52) | 0 (0-519) | Juxtatumor | 0.0078 |
| MDC | 198 (0-2264) | 45 (0-1226) | Tumor | 0.0083 |
| TGFa | 14 (0-209) | 9 (0-76) | Tumor | 0.0104 |
| IL-1RA | 1529 (17-10200) | 311 (0-10200) | Tumor | 0.0110 |
| Leptin | 12 (0-328) | 22 (0-426) | Juxtatumor | 0.0162 |
| MCSF | 2897 (634-27235) | 2124 (24-13266) | Tumor | 0.0173 |
| MIP-1b | 85 (4-517) | 45 (0-262) | Tumor | 0.0181 |
| CXCL10 | 527 (0-11000) | 106 (0-11000) | Tumor | 0.0200 |
| FGF-2 | 192 (29-1553) | 120 (0-501) | Tumor | 0.0233 |
| MCP-2 | 7 (0-151) | 13 (0-1037) | Juxtatumor | 0.0376 |
| CCL20 | 113 (0-8227) | 73 (0-547) | Tumor | 0.0496 |
| HGF | 2218 (115-8862) | 1195 (24-7529) | ns | 0.0621 |
| RANTES | 197 (4-5222) | 112 (0-3188) | ns | 0.0884 |
| TSLP | 0 (0-0) | 0 (0-13) | ns | 0.1250 |
| IL-8 | 11000 (3545-11000) | 11000 (2-11000) | ns | 0.1324 |
| LIF | 38 (0-731) | 75 (0-479) | ns | 0.1579 |
| IL-33 | 5 (0-135) | 15 (0-136) | ns | 0.2367 |
| I-309 | 0 (0-7) | 0 (0-3) | ns | 0.2789 |
| IL-23 | 0 (0-24) | 0 (0-24) | ns | 0.3750 |
| GRO | 12000 (236-12000) | 12000 (6-12000) | ns | 0.4634 |
| TPO | 0 (0-14) | 0 (0-22) | ns | 0.5000 |
| TNFb | 0 (0-2) | 0 (0-1) | ns | 0.6250 |
| G-CSF | 10500 (353-10500) | 10500 (0-10500) | ns | 0.6578 |
| MIP-1a | 207 (7-2100) | 193 (0-2100) | ns | 0.7152 |
| ENA-78 | 2212 (26-23000) | 2137 (0-23000) | ns | 0.8231 |
| CXCL6 | 65 (0-523) | 69 (0-2600) | ns | 0.8463 |
| CXCL7 | 1813 (144-7802) | 1447 (93-8201) | ns | 0.8815 |
| EGF | 4 (0-13) | 4 (0-27) | ns | 0.9809 |
| SDF-1 | 0 (0-77) | 0 (0-40) | ns | 1.0000 |
| IL-21 | 0 (0-0) | 0 (0-0) | ns | all values at 0 |
| IL-9 | 0 (0-0) | 0 (0-0) | ns | all values at 0 |

Cells highlighted in grey contain significant values at $p < 0.05$

Table S3 - Prognosis value of the 49 analytes measured in the tumor soluble microenvironment (Mann-Whitney)

| Analyte | Non-severe Median (min-max) | Severe Median (min-max) | p-value |
|---|---|---|---|
| MMP2 | 17432 (4155-51500) | 34839 (29414-51500) | 0.0074 |
| IL12(p70) | 1 (0-14) | 0 (0-2) | 0.0738 |
| EGF | 0 (0-13) | 7 (0-12) | 0.1422 |
| CCL20 | 82 (0-1160) | 303 (26-8227) | 0.1729 |
| MCP2 | 8 (0-151) | 0 (0-21) | 0.1934 |
| ENA78 | 2712 (26-23000) | 1468 (65-11471) | 0.2264 |
| CXCL9 | 52000 (61-52000) | 7350 (2415-52000) | 0.2286 |
| IL23 | 0 (0-24) | 0 (0-11) | 0.2501 |
| MCP3 | 0 (0-52) | 0 (0-0) | 0.2505 |
| IL1RA | 1137 (17-10200) | 2126 (421-10200) | 0.2958 |
| PAI1 | 19392 (1513-34000) | 22582 (12431-34000) | 0.3297 |
| CXCL6 | 85 (0-523) | 40 (5-394) | 0.3365 |
| IL1b | 748 (1-3519) | 1072 (88-5996) | 0.4122 |
| CXCL7 | 1348 (144-6613) | 2251 (535-7802) | 0.4325 |
| I309 | 0 (0-7) | 1 (0-2) | 0.4392 |
| TRAIL | 17 (0-238) | 20 (7-167) | 0.4679 |
| IL12(p40) | 0 (0-14) | 4 (0-24) | 0.5056 |
| TARC | 3 (0-87) | 7 (0-32) | 0.5351 |
| CXCL10 | 584 (0-11000) | 314 (168-1863) | 0.5588 |
| GRO | 10378 (236-12000) | 12000 (2966-12000) | 0.5810 |
| Resistin | 11045 (109-24500) | 8741 (413-24500) | 0.5851 |
| MMP9 | 10500 (783-10500) | 10500 (2806-10500) | 0.6027 |
| MMP1 | 21000 (7281-21000) | 21000 (21000-21000) | 0.6143 |
| TPO | 0 (0-14) | 0 (0-0) | 0.6143 |
| RANTES | 200 (4-5222) | 189 (98-1565) | 0.6385 |
| Leptin | 13 (0-226) | 6 (0-328) | 0.6957 |
| FGF2 | 159 (29-1553) | 250 (43-993) | 0.7149 |
| IL16 | 158 (18-2085) | 131 (35-1464) | 0.7411 |
| IL8 | 11000 (3545-11000) | 11000 (7858-11000) | 0.7421 |
| GCSF | 10500 (353-10500) | 10500 (1095-10500) | 0.7496 |
| TNFb | 0 (0-2) | 0 (0-1) | 0.7648 |
| IL15 | 5 (0-15) | 6 (2-17) | 0.7904 |
| IL33 | 3 (0-60) | 5 (0-135) | 0.8068 |
| LIF | 38 (0-731) | 67 (0-231) | 0.8593 |
| MIP1b | 87 (4-517) | 82 (31-177) | 0.8595 |
| MCP1 | 1103 (163-19500) | 1162 (269-12495) | 0.8734 |
| VEGF | 101 (0-2399) | 49 (26-1072) | 0.9154 |
| SDF1 | 0 (0-40) | 0 (0-77) | 0.9215 |
| MIP1a | 214 (7-2100) | 169 (60-889) | 0.9295 |
| MCSF | 2918 (634-12946) | 2639 (814-27235) | 0.9308 |
| SCF | 25 (0-93) | 21 (5-242) | 0.9435 |
| TNFa | 94 (1-2402) | 75 (29-1035) | 0.9584 |
| MDC | 184 (0-2264) | 219 (33-1050) | 0.9859 |
| TGFa | 16 (0-147) | 12 (5-209) | 0.9861 |
| HGF | 2218 (115-7258) | 2223 (363-8862) | 0.9861 |
| GMCSF | 1236 (0-10800) | 946 (753-10800) | 1.0000 |
| IL21 | 0 (0-0) | 0 (0-0) | 1.0000 |
| IL9 | 0 (0-0) | 0 (0-0) | 1.0000 |
| TSLP | 0 (0-0) | 0 (0-0) | 1.0000 |

Cells highlighted in grey contain significant values at $p < 0.05$

Table S4 - List of the 30 genes measured by RTqPCR

| Gene | Alias(es) | Included in the 18 gene signature |
|---|---|---|
| MMP1 | | no |
| MMP2 | | no |
| MMP9 | | no |
| CXCL10 | | no |
| CD3E | CD3 | no |
| FUT4 | CD15 | no |
| ICOSLG | ICOS-L | no |
| CD1C | | no |
| LAMP3 | | no |
| IL3RA | | no |
| CD8A | CD8 | no |
| PDCD1 | CD279, PD1 | no |
| CD274 | B7H1, PDL1, PDCD1L1 | yes |
| CCR7 | | yes |
| HLADRB | | yes |
| CCL5 | RANTES | yes |
| CD27 | TNFRSF7 | yes |
| CD276 | B7H3 | yes |
| CMKLR1 | | yes |
| CXCL9 | | yes |
| CXCR6 | | yes |
| HLA-DQA1 | | yes |
| HLA-E | | yes |
| IDO1 | IDO | yes |
| LAG3 | CD223 | yes |
| NKG7 | | yes |
| PDCD1LG2 | B7DC, PDL2 | yes |
| PSMB10 | LMP10 | yes |
| STAT1 | | yes |
| TIGIT | | yes |

Table S5 - Multivariate Cox proportional hazards Model 2, including n = 124 patients, and all clinical and biological data significant at $p < 0.05$ in univariate analysis

| Survival | Parameters | P value | HR (95% CI) |
|---|---|---|---|
| DSS | MMP2 | 0.034 | 1.168 (1.012-1.349) |
| | PNI | 0.020 | 2.599 (1.161-5.818) |
| OS | PNI | 0.010 | 2.198 (1.204-4.01) |
| | Stage | 0.019 | 2.646 (1.175-5.957) |
| DFS | MMP2 | 0.016 | 1.162 (1.028-1.312) |

Table S6 - Comparison of survivals in the prognostic groups defined by the Cox Model1

| | Prognostic groups | Log-rank | HR (Mantel-Haenszel) | | |
|---|---|---|---|---|---|
| | | P value | HR | Inf CI 95% | Sup CI 95% |
| DSS | MMP2 high / ENE- vs. MMP2 high / ENE+ | 0.0093 | 0.3417 | 0.1522 | 0.7671 |
| | MMP2 high / ENE- vs. MMP2 low / ENE- | 0.0022 | 3.228 | 1.524 | 6.834 |
| | MMP2 high / ENE- vs. MMP2 low / ENE+ | 0.6203 | 0.7928 | 0.3165 | 1.986 |
| | MMP2 high / ENE+ vs. MMP2 low / ENE- | <0.0001 | 21.49 | 7.226 | 63.94 |
| | MMP2 high / ENE+ vs. MMP2 low / ENE+ | 0.1851 | 1.795 | 0.7556 | 4.264 |
| | MMP2 low / ENE- vs. MMP2 low / ENE+ | 0.0016 | 0.1079 | 0.02715 | 0.4286 |
| DFS | MMP2 high / ENE- vs. MMP2 high / ENE+ | 0.0317 | 0.4281 | 0.1973 | 0.9285 |
| | MMP2 high / ENE- vs. MMP2 low / ENE- | 0.0893 | 1.771 | 0.916 | 3.426 |
| | MMP2 high / ENE- vs. MMP2 low / ENE+ | 0.6349 | 0.8029 | 0.3243 | 1.987 |
| | MMP2 high / ENE+ vs. MMP2 low / ENE- | 0.0002 | 5.539 | 2.236 | 13.72 |
| | MMP2 high / ENE+ vs. MMP2 low / ENE+ | 0.3634 | 1.497 | 0.6273 | 3.57 |
| | MMP2 low / ENE- vs. MMP2 low / ENE+ | 0.0705 | 0.3582 | 0.1177 | 1.09 |
| OS | MMP2 high/I or II vs. MMP2 high/III or more | 0.0402 | 0.5285 | 0.2873 | 0.972 |
| | MMP2 high/I or II vs. MMP2 low/I or II | 0.2129 | 1.886 | 0.6948 | 5.122 |
| | MMP2 high/I or II vs. MMP2 low/III or more | 0.653 | 6e-310 | 2e-322 | infinite |
| | MMP2 high/III or more vs. MMP2 low/I or II | 0.0004 | 2.8878 | 1.597 | 5.186 |
| | MMP2 high/III or more vs. MMP2 low/III or more | 0.0398 | 6e-310 | 2e-322 | infinite |
| | MMP2 low/I or II vs. MMP2 low/III or more | 0.0646 | 6e-310 | 2e-322 | infinite |

Inf: inferior. CI: confidence interval. Sup: infinite. Cells highlighted in grey contain significant values at $p < 0.05$

Table S7 - Clinical parameters according to *MMP2* RNA status

| Parameter | Percentage (n) | MMP2 Low (n=73) | MMP2 high (n=72) | p value (Fisher) | Odd Ratio [95%CI] |
|---|---|---|---|---|---|
| Gender | female | 40% (29) | 39% (28) | 1.0000 | |
| | male | 60% (44) | 61% (44) | 0.9506 | |
| Age | mean +/- SD | 63,21 +/- 13,68 | 63,35+/- 14,39 | | |
| Alcohol abuse (n=63, n=58) | absent | 59% (37) | 62% (36) | 0.7148 | |
| | present | 41% (26) | 38% (22) | | |
| Tobacco (n=70, n=67) | non smoker | 43% (30) | 43% (29) | 1.0000 | |
| | smoker | 57% (40) | 57% (38) | | |
| T stage | T1 or T2 | 40% (29) | 32% (23) | 0.3876 | |
| | T3 or T4 | 60% (44) | 68% (49) | | |
| N stage | N0 | 60% (44) | 42% (30) | 0.0310 | 2.11 [1.04; 4.35] |
| | N+ | 40% (29) | 58% (42) | | |
| Stage | I or II | 32% (23) | 24% (17) | 0.3536 | |
| | III or more | 68% (50) | 76% (55) | | |
| Differentiation | verrucous, well, basaloid | 75% (55) | 74% (53) | 0.8506 | |
| | moderate, poorly | 25% (18) | 26% (19) | | |
| Mitotic Index (n=63, n=56) | high | 54% (34) | 25% (14) | 0.0015 | 3.48 [1.51; 8.35] |
| | low / mid | 46% (29) | 75% (42) | | |
| Perineural invasion (n=63, n=62) | absent | 59% (37) | 37% (23) | 0.0200 | 2.40[1.12; 5.28] |
| | present | 41% (26) | 63% (39) | | |
| Vascular embols (n=65, n=61) | absent | 63% (41) | 59% (36) | 0.7157 | |
| | present | 37% (24) | 41% (25) | | |
| ENE | absent | 77% (56) | 69% (50) | 0.3536 | |
| | present | 23% (17) | 31% (22) | | |
| Margins | negative or close | 82% (60) | 83% (60) | 1.0000 | |
| | positive | 18% (13) | 17% (12) | | |
| HPV | negative | 93% (68) | 94% (68) | 1.0000 | |
| | positive | 7% (5) | 6% (4) | | |
| Adjuvant treatment | none | 41% (30) | 40% (29) | 0.9636 | |
| | RT | 41% (30) | 39% (28) | | |
| | RT + CT or Cetuximab | 18% (13) | 19% (14) | | |
| | curietherapy | 0% (0) | 1% (1) | | |
| Severity | low | 82% (60) | 65% (47) | 0.0241 | 2.44[1.07; 5.80] |
| | high | 18% (13) | 35% (25) | | |
| Recurrence | absent | 70% (51) | 51% (37) | 0.0398 | |
| | local | 30% (22) | 29% (21) | | |
| | regional | 14% (10) | 24% (17) | | |
| | metastatic | 7% (5) | 19% (14) | | |

Numbers in brackets beside clinical parameters indicate the number of patients for which the information was available. Cells highlighted in grey contain significant values at $p < 0.05$.

Table S8 - Primer sequences

| Primer Name | Primer Sequence 5' to 3' |
|---|---|
| D-ALB-U | GCTGTCATCTCTTGTGGGCTGT |
| D-ALB-L | ACTCATGGGAGCTGCTGGTTC |
| TBP-U | TGCACAGGAGCCAAGAGTGAA |
| TBP-L | CACATCACAGCTCCCCACCA |
| MMP1-U2 | GGCTTGAAGCTGCTTACGAATTT |
| MMP1-L2 | ACAGCCCAGTACTTATTCCCTTTGA |
| MMP2-U1 | ACTGCGGTTTTCTCGAATCCA |
| MMP2-L1 | GGTATCCATCGCCATGCTCC |
| MMP9-U1 | CGGCTTGCCCTGGTGCAGT |
| MMP9-L1 | CGTCCCGGGTGTAGAGTCTCTCG |
| CXCL10-U1 | CTGACTCTAAGTGGCATTCAAGGAG |
| CXCL10-L1 | GGTTGATTACTAATGCTGATGCAGG |
| CD3E-U2-Hs | AAGATGGTAATGAAGAAATGGGTGGT |
| CD3E-L2-Hs | TGAGGGCATGTCAATATTACTGTGGT |
| FUT4-U3-Hs | CTGCCATGGACCGTCTGTGT |
| FUT4-L3-Hs | CCCCAGCAAGCGTAGGTGA |
| CD274-U1-Hs | GCTGAATTGGTCATCCCAGAACTAC |
| CD274-L1-Hs | AAACGGAAGATGAATGTCAGTGCTAC |
| ICOSLG_U1_Hs | CTTCTGCAGCAGAACCTGACTGT |
| ICOSLG_L1_Hs | CGGTACTGACTGGATTCTCTGTGAT |
| CD1C-U1 | GACAATGCAGACGCATCCCA |
| CD1C-L1 | CAACTCGTCCAGCCATCCTGA |
| CCR7-U2 | GGGGAAACCAATGAAAAGCGT |
| CCR7-L2 | ATCTTGACACAGGCATACCTGGAA |
| LAMP3-U2 | ACCCGAAAATCCAACCTTCTGT |
| LAMP3-L2 | GTCAAATAGGCTCCCACTTCACTG |
| IL3RA-U1 | ATCGCAAATTTCGCTATGAGCTT |
| IL3RA-L1 | GGAGGTTCTGTCTCTGACCTGTTCT |
| HLA-class2-DRB-U2-Hs | TGCCAAGTGGAGCACCCAA |
| HLA-class2-DRB-L2-Hs | CAGATTCAGACCGTGCTCTCCAT |
| CCL5-U2 | GCCCACATCAAGGAGTATTTCTACA |
| CCL5-L2 | TTCGGGTGACAAAGACGACTG |
| CD27-U1-Hs | GTGCACCGAGTGTGATCCTCTT |
| CD27-L1-Hs | GGCCTCCAGCATCTCACTGAC |
| CD276-U1-Hs | AGGAGAATGCAGGAGCTGAGGA |
| CD276-L1-Hs | TCAGAGGCTGCAGGGCTGTC |
| CMKLR1-U2 | TCAACCTGGCAGTGGCAGAT |
| CMKLR1-L2 | CCCGAAAACCCAGTGGTAGTC |
| CXCL9-U2 | ATCCACCTACAATCCTTGAAAGAC |
| CXCL9-L2 | TCCATTCTTCAGTGTAGCAATGATTT |
| CXCR6-U1 | GGTTCAGCAGTTTCAATGACAGCA |
| CXCR6-L1 | CAGACCACAGACAAACACCACCAG |
| HLA-DQA1-U3 | CTACCGCTGCTACCAATGAGGTTC |

| | |
|---|---|
| HLA-DQA1-L3 | TGGGCTGACCCAGTGTCACG |
| HLA-E-U3 | GCTACTCTAAGGCTGAGTGGAGCGA |
| HLA-E-L3 | TTTACAAGCTGTGAGACTCAGACCCCT |
| IDO1-U1 | TGTTTCACCAAATCCACGATCAT |
| IDO1-L1 | CCTTCATACACCAGACCGTCTGAT |
| LAG3-U2-Hs | CCTTTCTCTGCTCCTTTTGGTGACT |
| LAG3-L2-Hs | AATCGTCTTGGTCGCCACTGTCT |
| NKG7-U1 | CCCCAGATCCAGACCTTCTTCTC |
| NKG7-L1 | CCAGGCTCAGGGCACCTGTA |
| PDCD1LG2-U1-Hs | TCCTGCTAATGTTGAGCCTGGAA |
| PDCD1LG2-L1-Hs | GTCACATTGCTGCCATGCTCTATTAT |
| PSMB10-U1 | CGCCCCCAAAATCTACTGCTG |
| PSMB10-L1 | TGGACGCCACCATCCGTGT |
| STAT1-U1 | AGCATGAAATCAAGAGCCTGGA |
| STAT1-L1 | ACCATTGGTCTCGTGTTCTCTGTT |
| TIGIT_Hs_U3 | CTCCCCTCGCCTCAGGAATGAT |
| TIGIT_Hs_L3 | CCGTGGTGGAGGAGAGGTGACA |
| CD8A-U3-Hs | CCGGTCTTCCTGCCAGCGAAG |
| CD8A-L3-Hs | GGCGCCGGTGTTGGTGGTC |
| PDCD1-U1-Hs | TCGTCTGGGCGGTGCTACAAC |
| PDCD1-L1-Hs | AGGGCCTGTCTGGGGAGTCTAAG |
